# Minimal net loss of vegetation via interior pond dynamics in an extensive unditched salt marsh

**DOI:** 10.1101/2020.07.20.212092

**Authors:** Joseph Smith, Michael Pellew

## Abstract

Ponds in salt marshes are often interpreted as a symptom of degradation, yet ponds can also be part of a cyclical process of pool formation, expansion, tidal breaching and vegetation recovery. Pond dynamics may be altered by accelerated sea level rise, with consequences for the long-term stability of ecosystems. We test the prediction that ponds are in dynamic equilibrium across one the largest expanses of unditched salt marsh in the Northeast USA by (1) examining change in pond and marsh area between 1970 and present and (2) by tracking individual pool dynamics across an 87-year time series. We found that net pond area has remained unchanged since 1970 because the amount of marsh conversion to ponds is equivalent to the amount pond recovery to marsh. The ratio of tidally-connected ponds is increasing relative to non-tidal ponds which suggests that some rates of change may be decoupling, which may be related to a decline in the rate of pond formation. A nuanced understanding of marsh pools needs to be incorporated into marsh condition assessments and establishment of restoration priorities so that ponds are not interpreted as evidence of degradation when they are exhibiting a recovery cycle. Unditched marshes around the world are a rare resource that remains essential for advancing scientific understanding and serving as reference sites for restoration of marshes altered by past management.

## Introduction

Around the world, efforts are ongoing to assess tidal marsh condition and resilience to accelerated sea level rise and insights from this work are guiding efforts to mitigate the impacts of climate change to extend the lifespan of marshes where possible (Kirwan and Megonigal 2013). At the same time, research continues to yield new insights into fundamental ecological and geomorphic processes in marshes. Hindering these efforts is the widespread prevalence of historic modifications of marshes, such as ditching and impoundment, that affect marsh condition and function in ways that potentially decrease resilience to sea level rise (Gedan et al. 2009). Tidal marshes without historic modifications are model systems for building upon fundamental knowledge of ecological processes and for understanding their inherent resilience to climate change and sea level rise. These insights can then be applied to improve management strategies in human-modified marshes that are intended to mitigate the impact of past modifications as well as the effects climate change.

Ponds in the marsh interior which occur in varying sizes, depths, densities and hydrologic settings, continue to be one of the least understood processes in salt marshes. Early descriptions of marsh ponds suggest that they are fundamental features of salt marshes (Miller and Egler 1950; Redfield 1972; Pethick 1974) and widespread ditching caused their extent to decline dramatically (Bourn and Cottam 1950; Lathrop et al. 2000; Adamowicz and Roman 2005). In a coastal resilience and restoration context, the limited understanding of pond dynamics in a variety of hydrologically altered and unaltered settings has led to ponds being broadly viewed as an indicator of degradation (Ganju et al. 2017), with ponds sometimes targeted with management actions intended to revegetate them (Wigand et al. 2017; Taylor et al. 2020). The distinction between marsh ponds as an inherent natural feature vs a symptom of degradation is critical to make so that management actions are directed appropriately. Ponds in some settings do represent degradation (Schepers et al. 2017) but they also can be a fundamental feature of marsh geomorphology (C. A. Wilson et al. 2014) and serve an important role as wildlife habitat (J. A. M. Smith and Niles 2016). This interpretation depends on a site’s management history as well as fundamental factors associated with elevation, tidal range and sediment flux (Mariotti 2016; Ganju et al. 2020).

A series of studies in New England (K. Wilson et al. 2009; K. Wilson et al. 2010; C. A. Wilson et al. 2014) demonstrated that ponds can be a source of resilience because they represent a dynamic process of pond formation and vegetation recovery. Ponds under this regime form on marsh above mean high water, expand over time and remain non-tidal until they are eventually breached by adjacent tidal creeks. Subsequently ponds experience rapid sedimentation and revegetate over a period of years. It is remarkable that a fundamental dynamic process influencing creek channel morphology, spatial variation in sedimentation (Mariotti et al. 2020) and plant community composition in salt marshes had been overlooked for so long. This may be because the majority of marshes have been so dramatically altered by ditching and impoundment that few places were left that still function in this way. To this point, Wilson (2014) had to account for the influence of ditching even where the pond process was still functioning. Because of former ditching and the limited study area extent, that study (C. A. Wilson et al. 2014) could pose the question, but could not fully answer, whether the ratio of marsh to pond area was in dynamic equilibrium.

In this study we document pond dynamics at a landscape scale across more than 5,000 Ha of salt marsh in southern New Jersey that has never been ditched or impounded. Salt marshes in the study area are predicted to be experiencing a “pond recovery regime” which entails pond formation, expansion, drainage, recovery (Mariotti 2016). We document salt marsh change over a 47-year time span to determine whether rates of pond formation and revegetation are at equilibrium or whether ponds represent a source of net marsh loss. To do this we partition patterns of marsh loss and gain along edges and the interior of both ditched and unditched areas. In addition, we track the formation, expansion and breaching dynamics of individual ponds in order to gain insights into whether change dynamics are exhibiting stable rates of change that are consistent with equilibrium dynamics.

This fundamental information on marsh change is needed to inform marsh vulnerability assessment and the development of management and restoration strategies. It is critical to understand how marshes are responding to sea level rise and other stressors in the absence of historic alterations so that the separate and interacting effects of sea level rise and historic alterations on marsh dynamics can be more clearly understood and management strategies can be better refined.

## Methods

### Study area

We defined our study area using National Wetlands Inventory (NWI) maps (Wilen and Bates 1995) classified as “Estuarine and Marine Wetland” or “Estuarine and Marine Deepwater” for New Jersey’s Atlantic coast tidal marshes that occur between barrier islands and the mainland. We constrained the study area to a 16,205 Ha region with extensive unditched areas from Cape May to Ocean City (Figure 1). As described below, we characterize all change within this footprint, but note that we do not estimate marsh expansion beyond this NWI footprint via marsh migration.

**Figure 1.**
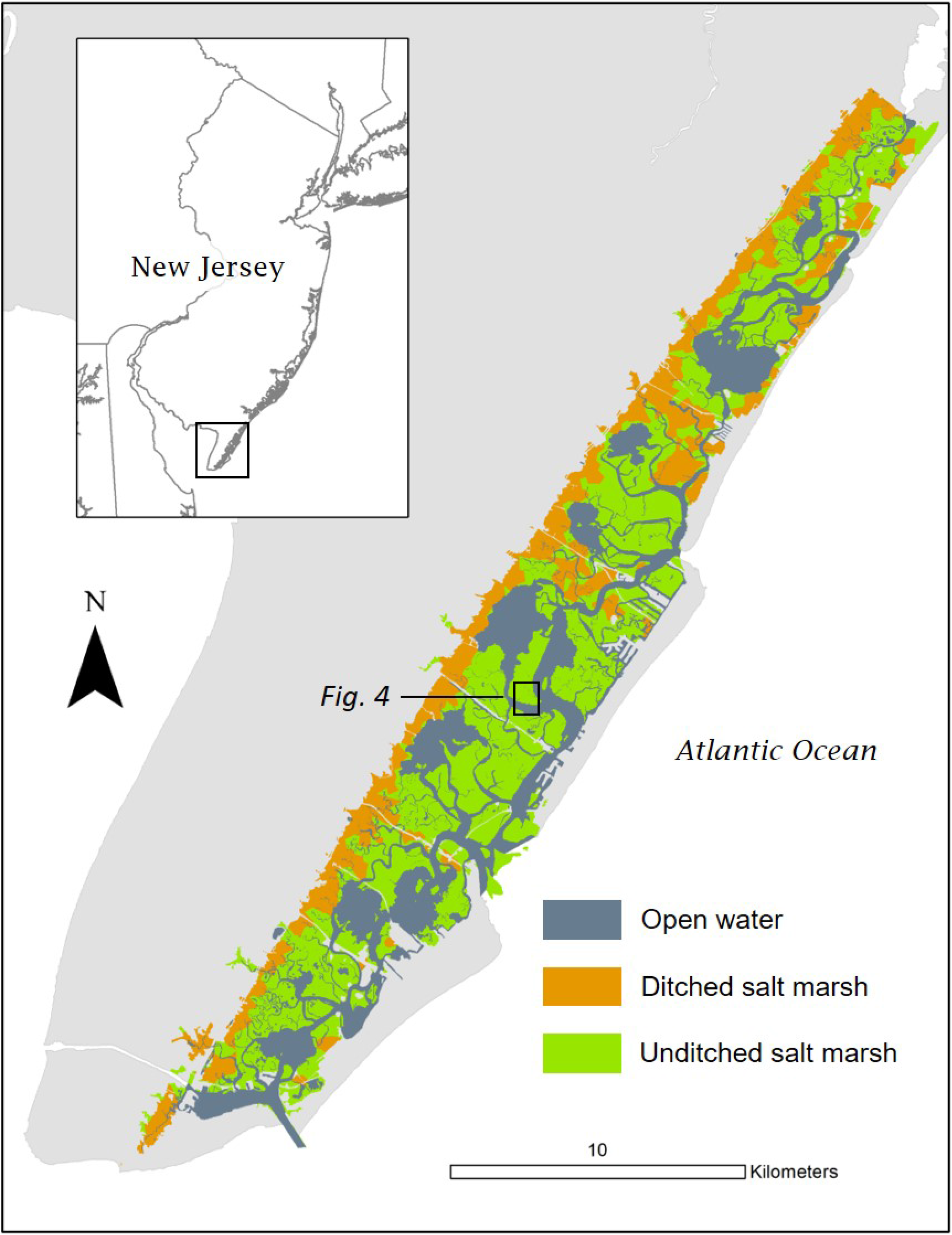
Study area defined by the National Wetlands Inventory areas classified as estuarine wetlands in Atlantic coast salt marshes from Cape May to Egg Harbor, NJ. Ditched and unditched areas were defined through classification of spatially-balanced random points with 100m buffers, where a marsh was considered to be ditched if a ditch occurred within the buffer area. For display purposes, points have been converted to Voronoi polygons. The inset rectangle locates the area depicted in Figure 4.

Previous hydrology work in the region concluded that the system is flood-dominated with suspended sediment concentrations ranging between 7-59 mg l ^−1^ depending on channel type and tidal stage (Ashley and Zeff 1988). Tidal range in the study area is 1.2-1.4m (Yang et al. 2008) and the 50-yr rate of relative sea level rise is 3mm/yr (Sallenger et al. 2012). Unditched salt marshes exhibit a sinuous channel morphology with 5 stream orders where 1^st^ order streams are dead end channels that terminate in the marsh interior (Zeff 1999). These areas also are characterized by tidal and non-tidal marsh ponds that occur on the marsh surface. Pond depths range from 3-60cm and size ranges from 1-55,000 m2 (Master 1992). Ponds in the region exhibit a cyclical dynamic where ponds form, expand, breach and revegetate over time (Mariotti 2016).

Ditching in the study area occurred in most cases prior to 1930 with activities focused primarily at the margins of uplands and barrier islands, whereas lagoonal marsh islands remained unditched.

### Historic and present marsh cover

We generated 1,560 spatially-balanced random points (Theobald et al. 2007) across the study area to estimate both historic and present marsh cover. At each point we classified land cover of the marsh surface for historic (1970) and present (2015-2017) time frames (Karl et al. 2014). This approach allows for estimation of cover at two points in time as well as the capacity to estimate patterns of change and stability among classes (Pontius et al. 2004).

We used 1970 NJ Wetlands Basemaps which are 1:24000 scale black and white aerial photos with tidal marsh plant species delineations (Brown 1978) superimposed on them. The maps were produced to support enforcement of the NJ Wetlands Act of 1970 (N.J.S.A. 13:9A-1, ET. SEQ). The 1970 baseline is notable in that from this point forward there was little direct filling and development of coastal wetlands in the study area. We also used 1:24,000 scale 1977 NJ tidelands Basemaps produced to support enforcement of the NJ Tidelands Act (N.J.S.A. 123) to verify classifications when there was uncertainty due to image quality or misalignment.

For the present time frame, we used 2017 true-color growing season imagery (USDA National Agriculture Imagery Program, 1 m pixel resolution aerial photography, maximum root mean square error (RMSE) 6m) to assign vegetation classifications and 2015 imagery (NJOIT OGIS orthoimagery, 0.3 m pixel resolution aerial photography) to confirm water cover classifications. Both images were collected during low tide.

Vegetation categories had unique spectral signatures that are easily distinguishable in true color aerial imagery during the growing season. We classified vegetated marsh into the following categories: tall *Spartina alterniflora*, short *S. alterniflora*, high marsh composed of *Spartina patens, Distichlis spicata* and/or Juncus gerardii, *Phragmites australis*, spoil placement sites and upland (we lump coastal shrubs such as *Iva fructescens* and *Baccharis halmifolia* into this category). We classified water cover as interior ponds without tidal creek connection (retaining water at low tide), interior ponds connected to tidal creeks, natural creeks, ditches, and open water of inlets and sounds. Several unique cover classes characterized marsh change between the two time intervals. These included marsh conversion to open water distinguished as “edge erosion” which documented marsh loss at bay edges, “creek expansion” which documented widening of tidal channels and “spoil recovery” where historic spoil placement had revegetated. Spoil classification includes only historic spoil disposal on marshes that occurred before wetlands laws outlawed this practice. Large confined disposal mounds that became the subsequent dredge disposal practice are not included in the analysis area. For each point we also noted its proximity to ditching and classified a point as occurring in ditched marsh if it was within 100 meters of a ditch. For analysis we delineated contiguous regions of sample points in unditched areas and excluded isolated points classified as unditched that typically occurred in upper watershed areas.

To estimate error around landcover percentages, we bootstrapped cover class frequency data with 1000 sampling iterations (Scheiner 1998). For categorical analyses of differences among landcover patterns based on point classification frequencies, we used fisher’s exact chi square test with Bonferroni correction for multiple comparisons (Rice 1989). We present results as area estimates based on percent cover of the study area delineated by the subsetted National Wetlands Inventory layer.

We used a recent tidal marsh mapping data set to conduct a partial validation of results (Correll et al. 2018) and used Lidar digital elevation models (NOAA Office for Coastal Management 2014) to describe relationships between elevation and vegetation patterns.

### Temporal variation in tidal breaching of ponds

We conducted a separate analysis to test whether the frequency of pond breaching by adjacent creeks is becoming more or less frequent over time. To do this we conducted a known-fate survival analysis (Kaplan and Meier 1958) of a subset of ponds over a span of 87 years (1930-2017). We selected ponds by taking a random sample of the points in unditched marsh that we used for landcover classification (n=80) and identifying the nearest non-tidal pond to the point. We divided the random points evenly into four cohorts corresponding with four time periods (1930, 1970, 1995 and 2006) to select ponds of varying age to include in the analysis. We delineated the size of each pond in GIS during the phase for which it was selected and then tracked each pond over the course of five time intervals (1930-56, 1957-1977, 1978-1995, 1995-2006 and 2006-2017) using a combination of imagery described below.

We used the 1970, 1977, 2015 and 2017 imagery described above along with imagery for 1931 (NJ Office of Information Technology (NJOIT), Office of Geographic Information Systems (OGIS)) which has a resolution of 2m per pixel and a root mean squared error (RMSE) of 11 m in coastal areas (J. A. M. Smith 2013). Imagery for 1995 is based on USGS digitial orthophotoquads with a 1m resolution and a maximum RMSE of 7m. Imagery for 2006 was from the USDA National Agriculture Imagery Program, with a pixel resolution of 1m and a maximum RMSE of 5m. We supplemental this imagery with additional time steps (1956 and 1963) available on historicaericals.com to determine the interval when ponds had breached.

Across the imagery series we noted patterns of change over time including pond expansion, breaching, revegetation and tidal channel formation. We analyzed pond breaching data with Program MARK (White and Burnham 1999) to implement the staggered entry form of this analysis (Pollock et al. 1989) to account for pond formation over the study period. We used the information-theoretic approach (Burnham and Anderson 2003) to evaluate variation in pond breaching by cohort (the time period during which a pond was selected), time period and pond age. Pond age was defined by the time interval in which a pond first appeared.

## Results

The total vegetated salt marsh in the study area was 8,945.3 Ha ± 388.9 SE in 1970 and 8,378.5 Ha ±372.8 SE in 2017, a 6.3% decrease. Approximately 60% of the marsh in the study area remains unditched, comprising a present-day vegetated marsh area of 5,5060 Ha which occurs primarily in lagoonal marsh areas disconnected from the mainland and barrier islands (Figure 1). Estimates of 2015-2017 salt marsh, open water (6,789.4 Ha) and pond area (502.2 Ha) was similar to Correll (2018) where remote sensing-derived area estimates were 8,240.3 Ha (salt marsh), 7093.5 Ha (open water) and 471.2 Ha (pond) using the same imagery. The predominant source of marsh loss (73% of total loss) was edge erosion, with 745.4 ± 80 Ha converting to open water (Figure 2).

**Figure 2.**
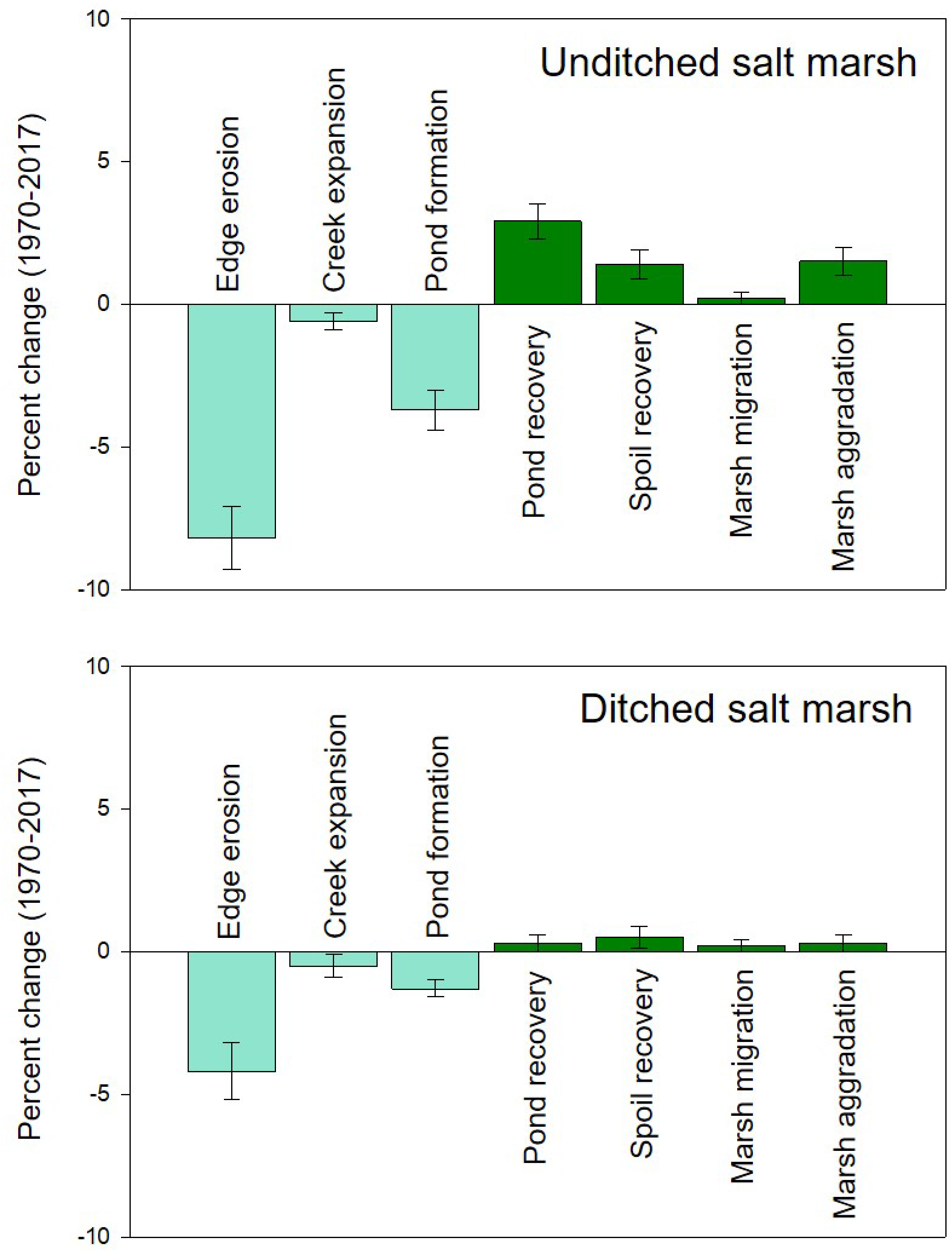
Patterns of change in unditched and ditched regions of our study area depicting losses and gains from the marsh edge and interior. Bars are estimated area in hectares ± SE.

Vegetated marsh is composed primarily of short-form *Spartina alterniflora* (64.2%) occupying areas above MHW, followed by tall-form *S. alterniflora* (32.1%) in areas between MTL and MHW (Figure 3). High marsh composed of *S. patens* and *Distichlis spicata* is a minor component (1.2%). Likewise, the invasive *Phragmites australis* is limited to the upland edge and former spoil placement areas (2.4%). The primary pattern of swap among vegetation classes from 1970 to 2017 (Table S1) was from short alterniflora to tall alterniflora (308.7 Ha ± 64.8 SE). Conversion from short to tall alterniflora was related to edge erosion and channel expansion (45% of observations), pond recovery (38.7%) and proximity to the northern edge of road beds crossing marshes (within 25m, 16.1%), where wrack accumulation from nor’easter storms causes recurring vegetation disturbance. In addition, high marsh converted to *Spartina alterniflora* at a high rate, resulting in a 53% reduction in high marsh area during the period of study (from 243.1 ± 48.6 to 113.4 Ha ± 32.4 SE). Net vegetated marsh loss in unditched marsh was 6.3% (Figure 2, loss: −8.2% edge erosion, −3.5% pond formation, −0.6% channel expansion; gains: 0.2% marsh migration, 1.4% spoil recovery, 2.9% pond recovery, 1.5% marsh aggradation). Net vegetated marsh loss in ditched marsh was 3.3% (Figure 2, loss: −4.2% edge erosion, −1.3% pond formation, −0.5% channel expansion); gains: 1.6% marsh migration, 0.5% spoil recovery, 0.3% pond recovery, 0.3% marsh aggradation). Patterns of vegetated marsh gain and loss fell into two broad categories: edge and interior processes. Edge processes encompass losses via edge erosion and channel expansion as well as gains via marsh aggradation and marsh migration into uplands. Interior processes include marsh loss via pond formation, gains via pond revegetation and pond transition from a non-tidal to tidal hydrology (Figure 4). The ratio of marsh to interior open water (tidal and non-tidal ponds) is 9.7% in unditched areas and 2.4% in ditched areas. These ratios did not change significantly between 1970 and 2017 in either marsh type (unditched Χ^2^=0.14, DF=1, p= 0.71, ditched Χ^2^=0.21, DF=1 p=0.65).

**Figure 3.**
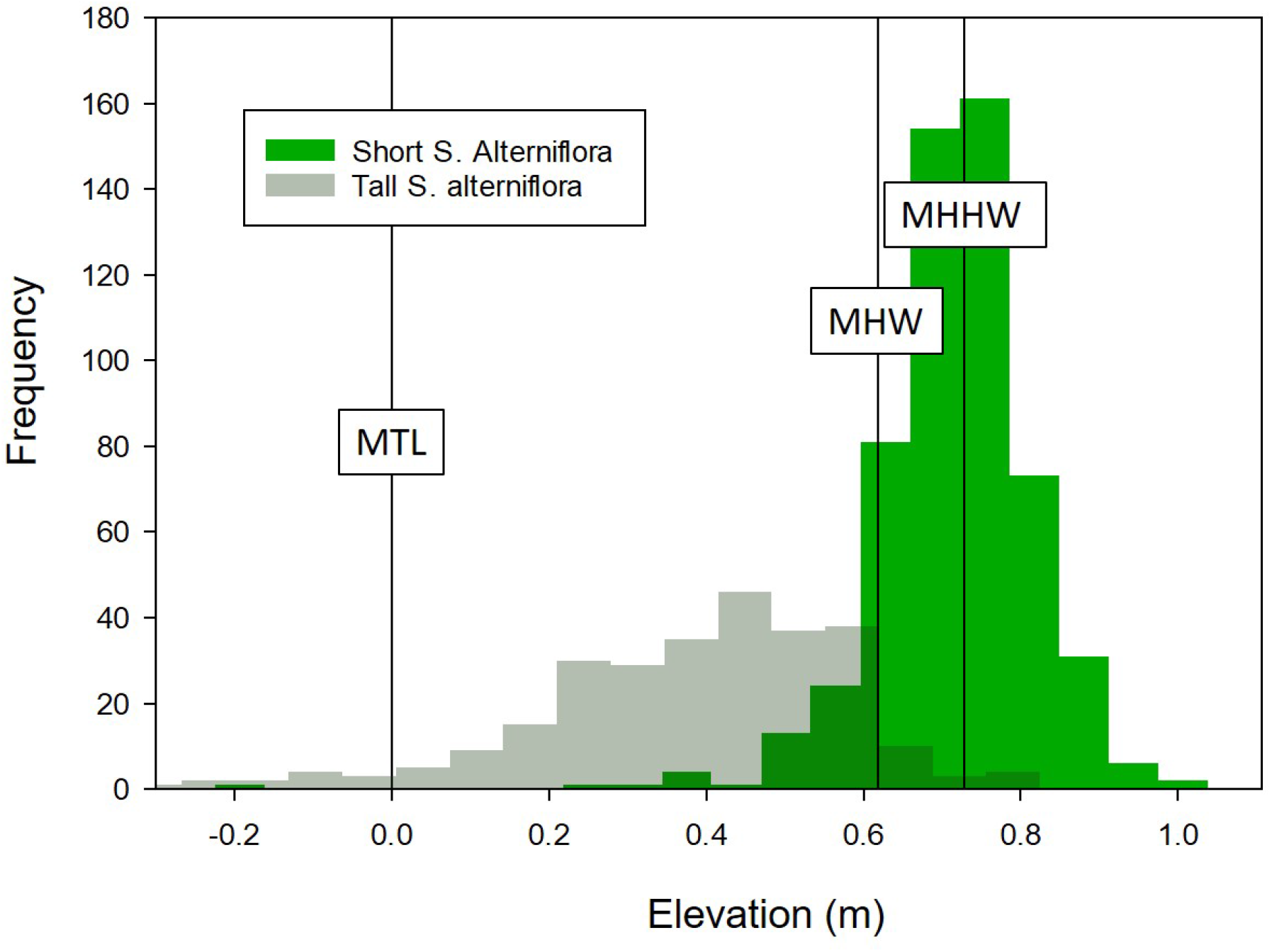
Elevation-vegetation relationships for the two dominant vegetation classes in the study area. The tall form of Spartina alterniflora occupied elevation between MTL and MHW while the short form of this species occupied higher elevations where ponds typically form. Elevation is based on a 2014 USGS LiDAR digital elevation model.

**Figure 4.**
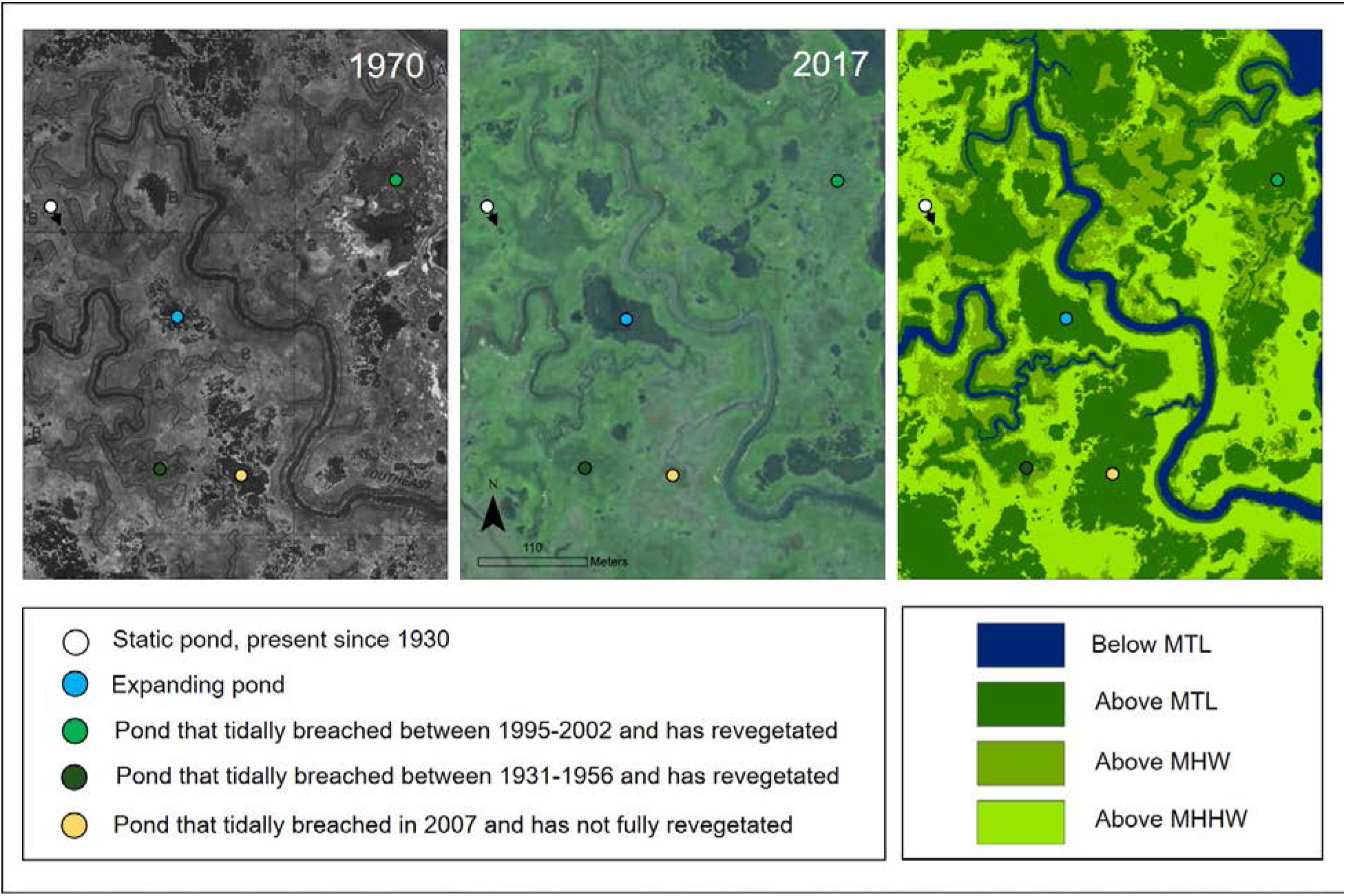
Typical undtiched marsh area depicted in 1970 and 2017, along with LiDAR digitial elevation model binned to depict tidal datums. Examples of different aspects of pond dynamics are noted with colored points.

Unditched marsh showed greater overall dynamics for both edge and interior processes (Figure 2). In unditched marsh, edge dynamics (vegetation increase or decrease at marsh edge) were evident for 10.6% of observations, while in ditched marsh this value was 5.7% (Χ^2^=7.7, DF=1, p= 0.0054). This difference was due primarily to distribution of the two marsh types in the landscape. Unditched lagoonal marshes experienced greater amounts of edge erosion due to their exposure to large open water bodies (Mariotti and Fagherazzi 2013) while ditched marshes adjacent to uplands were buffered from such erosion. Dynamics in the marsh interior comprising exchange between open water and vegetated marsh was 6.4% for unditched while in ditched marsh such dynamics were evident in only 1.7% of observations. (Χ^2^=14.1, DF=1, p= 0.0002).

### Pond dynamics in unditched marsh

In unditched marshes, the estimated areas of pond formation/expansion (228.6 Ha ± 32.7) and pond revegetation (189.8 ± 39.3 SE) since 1970 were not significantly different (Figure 5, Χ^2^= 0.6, DF=1, p= 0.54). This indicates that due to the cyclic dynamics of pond formation and recovery, there was little net change to vegetated marsh due to pond formation and expansion. Marsh that converted to ponds was short *Spartina alterniflora* while ponds that revegetated were composed of tall S. alterniflora, which reflects lower elevations of recovering ponds.

**Figure 5.**
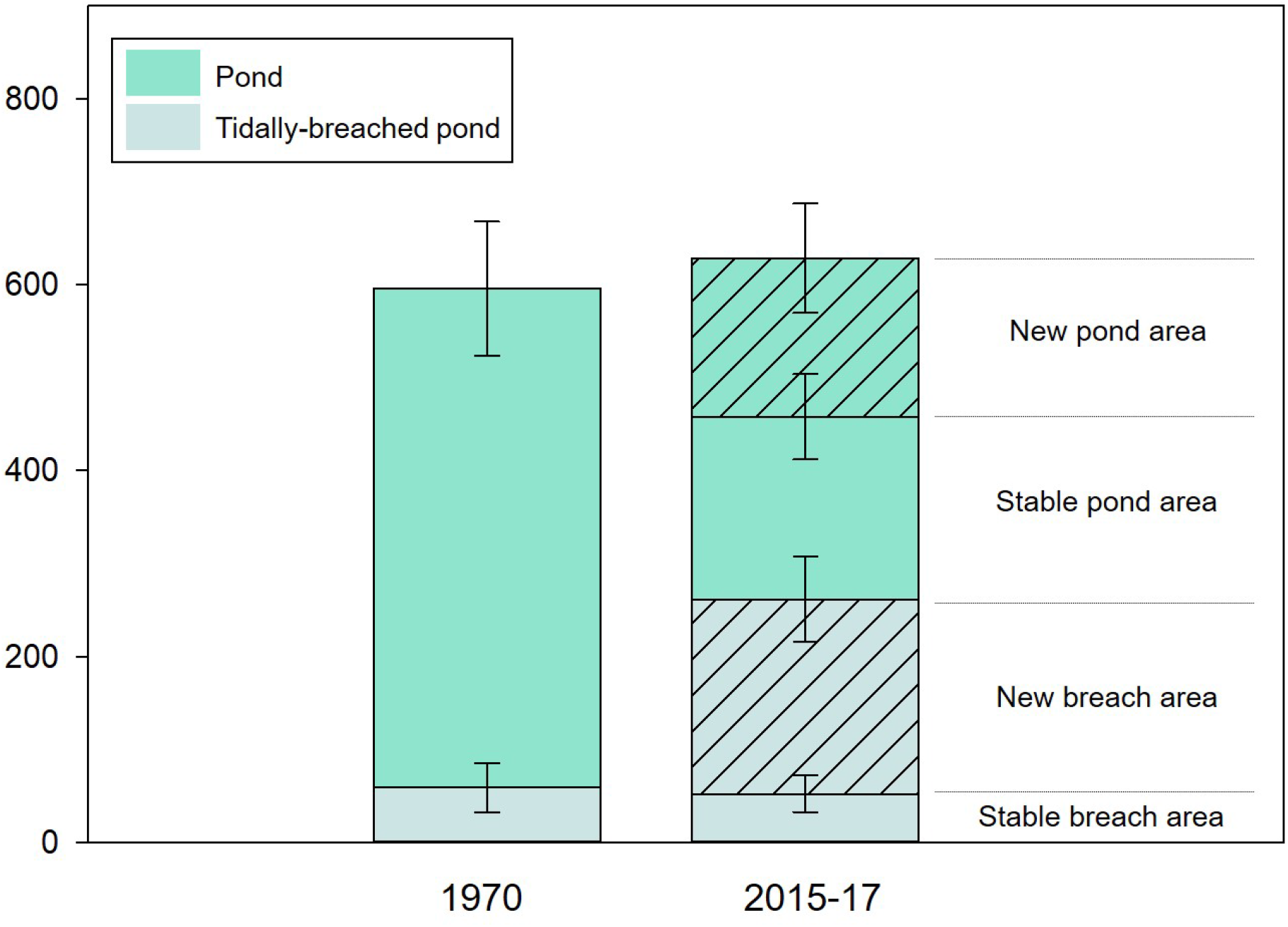
Patterns of change in pond area between 1970 and 2017, indicating no significant difference in overall pond area. The proportion of tidal to non-tidal ponds increased significantly. The fractions of retained vs new open water are noted for the latter interval. The formation of new open water was offset by pond revegetation in other areas (see Figure 2). Bars are estimated area in hectares ± SE.

A further examination of patterns of stability and change of interior open water features revealed that for non-tidal ponds (not connected to tidal creeks), approximately half of the area occupied by ponds in 2015-17 was also occupied in 1970 while the remaining half has formed since 1970 (Figure 5, via expansion of existing ponds (86% of observations) or formation of new ponds (14%). Non-tidal ponds occurred almost exclusively in areas short-form S. alterniflora on marsh that is above MHW. For ponds that have breached and are connected to tidal creeks, area has increased significantly from 58.9 ± 26.2 Ha in 1970 to 261.8 ± 45.8 Ha in 2015-2017 (Χ^2^= 12.8, DF=1, p= 0.0005, Figure 3). Nontidal pond area correspondingly decreased from 536.6 ± 72 Ha in 1970 to 379.5 ± 58.9 Ha, but this difference was not statistically significant (Χ^2^= 3.0, DF=1, p= 0.083).

### Dynamics of individual ponds

We examined the history of 75 ponds across 4 time periods ranging from 1930 to 2017 (5 of the original sample of 80 were omitted due to proximity to human disturbance). Ponds ranged in size from 0.004 to 3.6 Ha. 65% of the ponds were present since 1930, with 31% appearing between 1957-77 and the remaining 4% appearing between 1978-95. All of the 20 randomly-selected ponds from 2006 images appeared prior to 1996.

The majority of ponds were dynamic (88%, i.e. expanding over time) with 29 of 66 becoming breached by tides during the study. Dynamic pond size did not vary significantly across the four time periods (Wilcoxon Χ^2^=6.98, DF=3, p= 0.073). Unexpectedly, a subset of ponds (9 of 75, 12%) were static in size and present from the start to the end of the study period. Static ponds were smaller (0.015 Ha ± 0.0005 SE, n=10) than dynamic ponds (0.42 Ha ± 0.069 SE, n=65).

The majority of the 29 ponds that became tidally breached throughout the study period showed a steady progression of revegetation over time (Figure 6) and the formation of small tidal channels that bisect the former pond area (Figure 4). The ponds that breached in the early to mid-20^th^ century have experienced complete revegetation, with well-defined tidal channels evident in all cases, while more recently breached ponds (since 2006) are just beginning to revegetate (Figure 6) and experience channel formation (3 of 9 breached ponds with channels).

**Figure 6.**
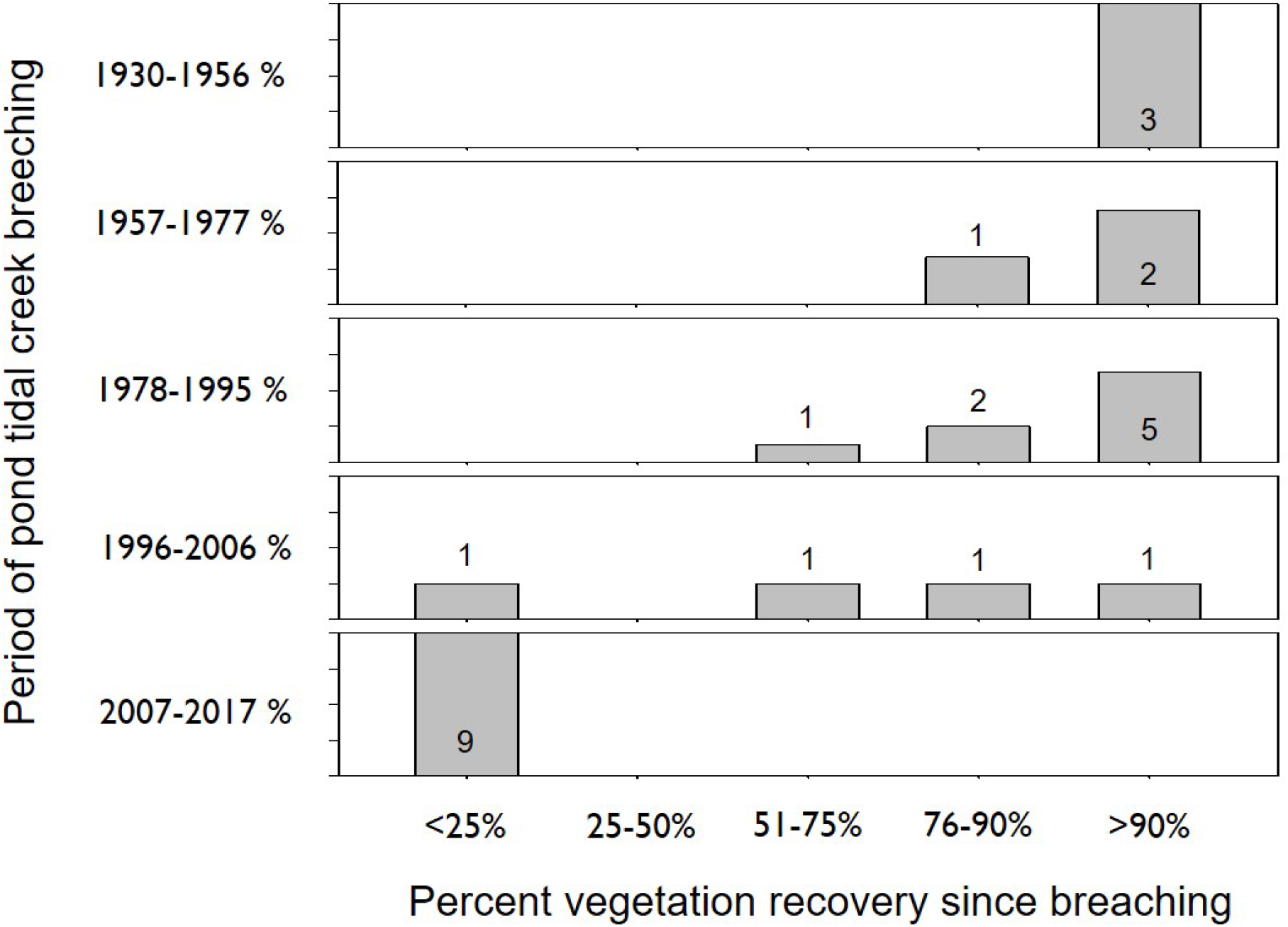
Revegetation progress of randomly-selected ponds that breached between 1930 and 2017 divided into five time intervals. Bars represent the percent of ponds in each revegetation category and numbers displayed with bars are raw count of ponds.

The probability of pond breaching varied according to the three factors we examined: by time interval, by cohort (the period when ponds were selected) and by pond age. The probability of breaching increased through the five time steps analyzed (Figure 7 top panel) while pond cohorts selected from earlier aerial imagery were more likely to breach over the course of the study than ponds selected from more recent imagery (Figure 7 top panel). These two patterns reveal opposing trends that that can be explained by an increasing prevalence of older ponds which have an increasing likelihood of breaching with age (Figure 7, bottom panel). Pond age was the best supported predictor of breaching probability, with time and cohort models (as well as a null model of no variation in breaching probability) having a ΔAICc difference of >10 compared with the pond age model.

**Figure 7.**
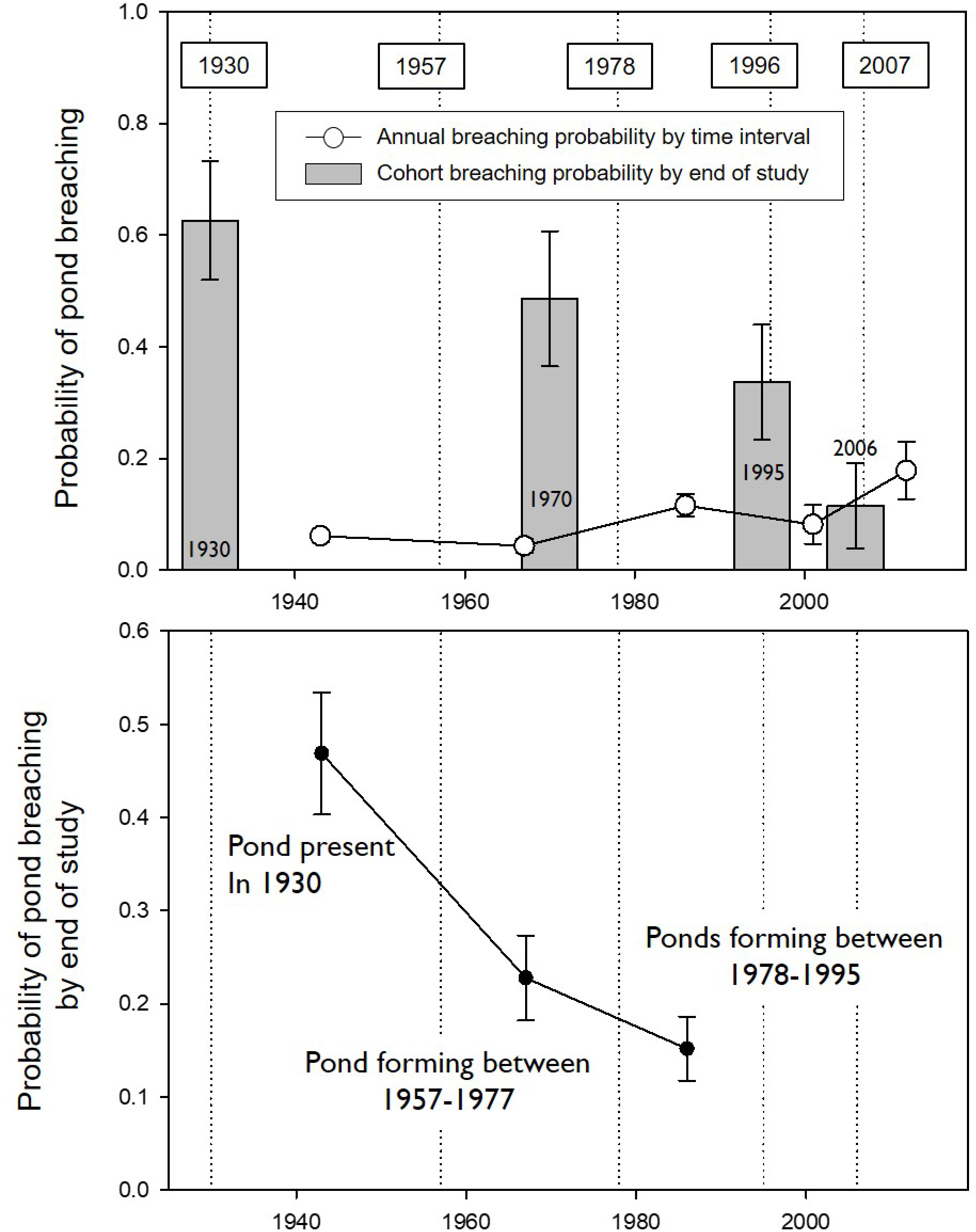
Time-to-event analysis of tidal breaching of ponds based on randomly selected ponds (n=75) drawn from aerial photos at four time phases (1930, 1970, 1995, 2006) and tracked across 5 time intervals between 1930 and 2017. Absolute pond breaching rate increased over time (top panel, line graph) although randomly-selected ponds from more recent time phases had a lower probability of breaching (top panel, bars). This pattern is best explained by the increasing overall age of ponds, with ponds becoming more likely to breach as they age (bottom panel).

## Discussion

The detailed dynamics of salt marsh change described here across thousands of hectares of unditched salt marsh validate the prediction that ponds in the interior of these marshes are not a significant net contributor to marsh loss (Mariotti 2016). Overall, the vegetated marsh area decreased by 6.3%, which is moderate compared to other estimates of salt marsh change in the region (Watson et al. 2016; Campbell and Wang 2019). The primary source of loss is edge erosion, which is caused in many parts of the study area by boat wake and wind-driven waves in coastal bays (Mariotti and Fagherazzi 2013) rather than sea level rise. This process likely plays an important role in providing material for marsh accretion and for pond recovery (Hopkinson et al. 2018). Tidal channel expansion plays a relatively minor role in marsh loss and did not vary between ditched and unditched marsh, which is consistent with other measurements for this study area (Robinson 2018).

Compared with unditched marsh, ditched marshes experienced lower overall change dynamics, primarily because ponds are less frequent in these areas. One of the purposes of ditching was to eliminate interior open water on marshes (J. B. Smith 1907) and the pattern of decreased pond area in ditched marshes has been documented elsewhere (Bourn and Cottam 1950; Lathrop et al. 2000; Adamowicz 2002). The lack of ponds and their associated dynamics may ultimately translate to lower resilience to sea level rise.

The pond cycle is inherently resilient because marsh loss and gain are in constant interchange and a complex sediment delivery system is in place that brings sediment directly to the marsh interior (C. A. Wilson et al. 2014). Small creeks that dead end into settling basins that were formerly ponds are ideally suited for concentrating and settling out fine sediment (Zeff 1988). Over longer time periods, regions of the marsh experiencing higher and lower rates of sediment accumulation alternate as a result of the pond cycle (Mariotti et al. 2020). Conversely, ditched marshes tend to have elevation deficits and are sediment-starved in the interior (LeMay 2007), with the ditches themselves becoming sediment sinks (Corman et al. 2012). Ditched marshes in this study did not experience greater amounts of vegetated area loss compared with unditched marsh, which may be related to tidal range. The consequences of altered hydrology and sedimentation patterns likely become greater as tidal range becomes narrower (Kearney and Turner 2016). For example, in microtidal ditched marshes, there is a pervasive pattern of permanent open water formation between ditches (Watson et al. 2016) that is not evident in our study area. In addition, with the pond cycle absent in ditched marshes, the resulting lack of heterogeneity in hydrologic features, elevation and vegetation yields a less diverse array of habitats for fish and wildlife (J. A. M. Smith and Niles 2016).

Given the fundamental importance of the pond cycle in dictating sedimentation patterns, tidal channel network morphology, elevation, vegetation patterns and habitat availability, it is critical to understand the mechanisms initiating and perpetuating pond dynamics. This knowledge is important for both understanding the response of these landscape to climate change, but also for informing the design of projects intended to restore natural hydrology to ditched and impounded marshes. There are a range of interacting factors that dictate rates of pond formation, expansion, tidal breaching, dead-end tidal channel formation, revegetation to marsh below MHW and subsequent dead-end channel infill and marsh accretion that bring the cycle back to its beginning. Each of these processes occur at an inherent rate that can be independent or dependent on other processes. All must be synchronized in order to arrive at an equilibrium distribution of geomorphic features on the landscape.

Patterns in our data suggest that some of these processes may be coming out of sync. For example, the rate of pond formation is potentially slowing because most of the new pond area formed since 1970 occurred via expansion of existing ponds (86%) and similarly we did not encounter any ponds that formed since 1995 in our random selection of ponds in 1995 and 2006. It is difficult to know if this pattern deviates from an equilibrium rate, but further insight can be gained from the pond survival analysis. Here we saw a pattern of changing age distribution of ponds over time. If the age-distribution of ponds was constant, as would be predicted of an equilibrium state, we would observe similar rates of breaching over time. We instead saw an increasing rate of breaching over time that was due to an increasing prevalence of older ponds. These older ponds have longer periods to expand in size and so become increasingly likely to intersect with adjacent tidal creeks. The increased ratio of breached to unbreached ponds we documented may be due in part to a slowing of pond formation, but a decline in the revegetation rate of breached ponds cannot be ruled out as a contributing factor.

Pond formation is probably the least-understood part of the pond cycle. The many proposed explanations include waterlogging and reduced sedimentation (Redfield 1972) as well as various forms of vegetation disturbance by ice (Dionne 1968), wrack (Harshberger 1916), or burrowing/grazing organisms such as crabs (Escapa et al. 2015). Fundamentally, the process associated with expanding ponds within our study area involves a disturbance to vegetation that creates conditions hostile to subsequent vegetation regrowth long enough for the kernel of a pool to form (Perillo 2019).

The static ponds we observed in our study area that experienced little or no expansion and were present for the entire 87-year course of study likely have a different origin than the expanding pools. These are akin to the “potholes” described in a Connecticut marsh (Miller and Egler 1950). Pond genesis is often described in terms of primary origins (existing before the marsh formed) or secondary origins (forming on established marsh). Expanding ponds are clearly of secondary origin, but we cannot rule out the possibility that static ponds are of primary origin.

At this stage, the impacts to marsh resilience from past management are widely recognized but are difficult to disentangle from the impacts of sea level rise (Kirwan and Megonigal 2013). This is the case because of the ubiquity of direct management of marsh hydrology for a range of human uses (Silliman et al. 2009). Given this reality, marshes that have not been subject to physical manipulations (e.g. ditching and impoundment) serve as an essential resource for both building upon basic knowledge of ecogeomorphic processes and for understanding marsh resilience to sea level rise apart from the impact of past management. Both of these opportunities for learning then lead directly to developing better solutions for restoring and managing the vast areas of marshes that have been degraded by past manipulations.

Although it is clear from modelling predictions and the empirical results of this study that interior open water is not always the result of sea level rise-driven degradation, there are still calls for addressing these features as such (Taylor et al. 2020). Such conclusions generally stem from interpretations of marsh features that (1) do not critically examine whether interior open water is an artefact of past management, as is seen in ditched and formerly impounded marsh (J. A. M. Smith et al. 2017), and (2) do not consider of the combination of marsh attributes that determine whether ponds represent marsh drowning, pond collapse or pond recovery regimes (Mariotti 2016). When these factors are considered, a clearer perspective on marsh trajectory and resilience can be gained and in turn, a clearer agenda for marsh conservation and restoration can be developed.

---

**Table S1.**
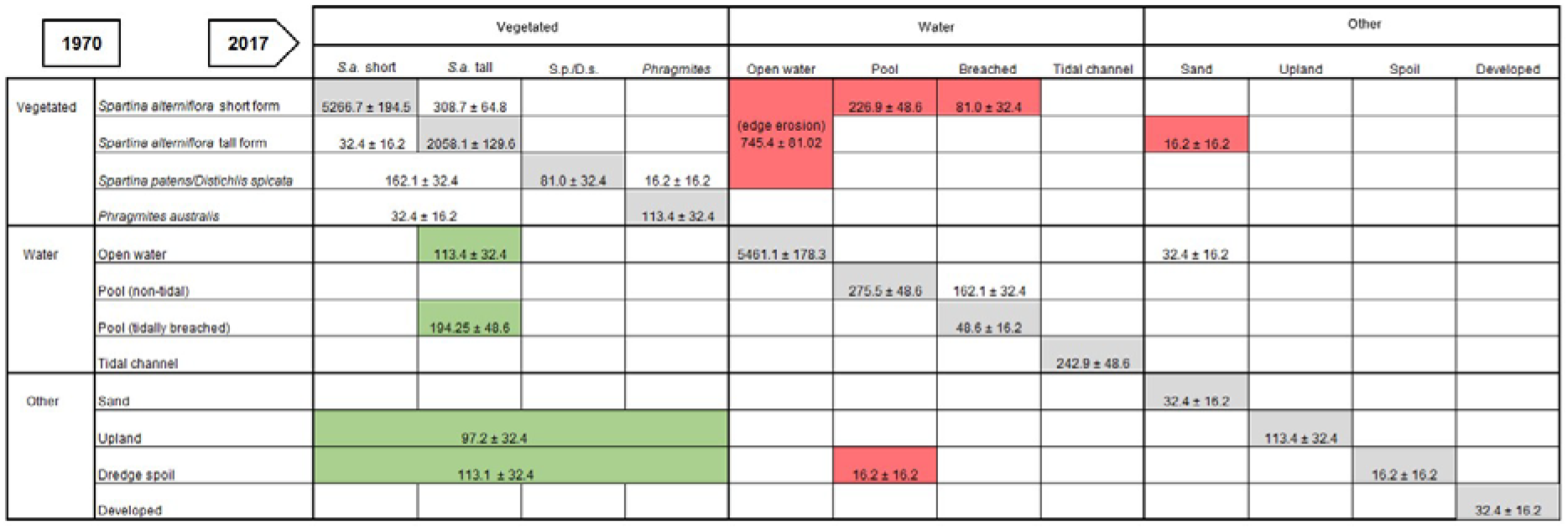
Patterns of stability and change between cover classes between 1970 and 2017. The area of each category that remained stable is highlighted in gray. Patterns of change that resulted in an increase in vegetated marsh are highlighted in green and those that resulted in a decrease are highlighted in red.

## Literature Cited

Adamowicz, Susan C. 2002. New England Salt Marsh Pools: Analysis of Geomorphic and Geographic Parameters, Macrophyte Distribution and Nekton Use.

Adamowicz, Susan C., and Charles T. Roman. 2005. New England salt marsh pools: A quantitative analysis of geomorphic and geographic features. Wetlands 25: 279–288.

Ashley, Gail M., and Marjorie L. Zeff. 1988. Tidal channel classification for a low-mesotidal salt marsh. Marine Geology 82. Elsevier: 17–32.

Bourn, Warren Scudder, and Clarence Cottam. 1950. Some Biological Effects of Ditching Tidewater Marshes. Washington D.C.: U.S. Fish and Wildlife Service.

Brown, W. W. 1978. Wetland mapping in New Jersey and New York. Photogrammetric Engineering and Remote Sensing 44: 303–314.

Burnham, Kenneth P., and David Anderson. 2003. Model Selection and Multi-Model Inference. 2nd ed. Springer.

Campbell, Anthony, and Yeqiao Wang. 2019. High spatial resolution remote sensing for salt marsh mapping and change analysis at fire island national seashore. Remote Sensing 11. Multidisciplinary Digital Publishing Institute: 1107.

Corman, Sarah S., Charles T. Roman, John W. King, and Peter G. Appleby. 2012. Salt Marsh Mosquito-Control Ditches: Sedimentation, Landscape Change, and Restoration Implications. Journal of Coastal Research: 874–880.

Correll, Maureen D., Wouter Hantson, Thomas P. Hodgman, Brittany B. Cline, Chris S. Elphick, W. Gregory Shriver, Elizabeth L. Tymkiw, and Brian J. Olsen. 2018. Fine-Scale Mapping of Coastal Plant Communities in the Northeastern USA. Wetlands: 1–12. https://doi.org/10.1007/s13157-018-1028-3.

Dionne, Jean-Claude. 1968. Action of shore ice on the tidal flats of the St. Lawrence estuary. Atlantic Geology.

Escapa, Mauricio, Gerardo ME Perillo, and Oscar Iribarne. 2015. Biogeomorphically driven salt pan formation in Sarcocornia-dominated salt-marshes. Geomorphology 228. Elsevier: 147–157.

Ferrarin, Christian, Fantina Madricardo, Federica Rizzetto, William Mc Kiver, Debora Bellafiore, Georg Umgiesser, Aleksandra Kruss, Luca Zaggia, Federica Foglini, and Alessandro Ceregato. 2018. Geomorphology of scour holes at tidal channel confluences. Journal of Geophysical Research: Earth Surface 123. Wiley Online Library: 1386–1406.

Ganju, Neil K., Zafer Defne, and Sergio Fagherazzi. 2020. Are Elevation and Open-Water Conversion of Salt Marshes Connected? Geophysical Research Letters 47: e2019GL086703. https://doi.org/10.1029/2019GL086703.

Ganju, Neil K., Zafer Defne, Matthew L. Kirwan, Sergio Fagherazzi, Andrea D’Alpaos, and Luca Carniello. 2017. Spatially integrative metrics reveal hidden vulnerability of microtidal salt marshes. Nature Communications 8: 14156. https://doi.org/10.1038/ncomms14156.

Gedan, K. B, B. R. Silliman, and M. D. Bertness. 2009. Centuries of human-driven change in salt marsh ecosystems. Annual Review of Marine Science 1: 117–141.

Goudie, Alice. 2013. Characterising the distribution and morphology of creeks and pans on salt marshes in England and Wales using Google Earth. Estuarine, Coastal and Shelf Science 129. Elsevier: 112–123.

Harshberger, John W. 1916. The origin and vegetation of salt marsh pools. Proceedings of the American Philosophical Society: 481–484.

Hopkinson, Charles S., James T. Morris, Sergio Fagherazzi, Wilfred M. Wollheim, and Peter A. Raymond. 2018. Lateral marsh edge erosion as a source of sediments for vertical marsh accretion. Journal of Geophysical Research: Biogeosciences 123. Wiley Online Library: 2444–2465.

Kaplan, E. L, and P. Meier. 1958. Nonparametric estimation from incomplete observations. Journal of the American Statistical Association 53: 457–481.

Karl, Jason W., Jeffrey K. Gillan, Nichole N. Barger, Jeffrey E. Herrick, and Michael C. Duniway. 2014. Interpretation of high-resolution imagery for detecting vegetation cover composition change after fuels reduction treatments in woodlands. Ecological indicators 45. Elsevier: 570–578.

Kearney, Michael S., and R. Eugene Turner. 2016. Microtidal Marshes: Can These Widespread and Fragile Marshes Survive Increasing Climate–Sea Level Variability and Human Action? Journal of Coastal Research 32. Allen Press: 686–699. https://doi.org/10.2112/JCOASTRES-D-15-00069.1.

Kirwan, Matthew L., and J. Patrick Megonigal. 2013. Tidal wetland stability in the face of human impacts and sea-level rise. Nature 504: 53–60.

Lathrop, R.G., M.B. Cole, and R.D. Showalter. 2000. Quantifying the habitat structure and spatial pattern of New Jersey (U.S.A.) salt marshes under different management regimes. Wetlands Ecology and Management 8: 163–172.

LeMay, Lynsey E. 2007. The Impact of Drainage Ditches on Salt Marsh Flow Patterns, Sedimentation and Morphology: Rowley River, Massachusetts. M.S. thesis, College of William and Mary.

Linthurst, Rick A., and Ernest D. Seneca. 1980. The effects of standing water and drainage potential on the Spartina Alterniflora-substrate complex in a North Carolina salt marsh. Estuarine and Coastal Marine Science 11: 41–52. https://doi.org/10.1016/S0302-3524(80)80028-4.

Mariotti, and Sergio Fagherazzi. 2013. Critical width of tidal flats triggers marsh collapse in the absence of sea-level rise. Proceedings of the National Academy of Sciences of the United States of America 110: 5353–5356. https://doi.org/10.1073/pnas.1219600110.

Mariotti, G. 2016. Revisiting salt marsh resilience to sea level rise: Are ponds responsible for permanent land loss? Journal of Geophysical Research: Earth Surface 121: 1391–1407.

Mariotti, G., A. C. Spivak, S. Y. Luk, G. Ceccherini, M. Tyrrell, and M. Eagle Gonneea. 2020. Modeling the spatial dynamics of marsh ponds in New England salt marshes. Geomorphology 365: 107262. https://doi.org/10.1016/j.geomorph.2020.107262.

Master, Terry L. 1992. Composition, structure, and dynamics of mixed-species foraging aggregations in a southern New Jersey salt marsh. Colonial Waterbirds: 66–74.

McKee, Karen L., Irving A. Mendelssohn, and Michael D. Materne. 2004. Acute salt marsh dieback in the Mississippi River deltaic plain: a drought-induced phenomenon? Global Ecology and Biogeography 13: 65–73. https://doi.org/10.1111/j.1466-882X.2004.00075.x.

Miller, William R., and Frank E. Egler. 1950. Vegetation of the Wequetequock-Pawcatuck Tidal-Marshes, Connecticut. Ecological Monographs 20: 143–172.

NOAA Office for Coastal Management. 2014. NOAA NGS Topobathy Lidar DEM: Post-Sandy (SC to NY).NOAA National Centers for Environmental Information.

Perillo, Gerardo ME. 2019. Geomorphology of Tidal Courses and Depressions. In Coastal Wetlands, 221–261. Elsevier.

Pethick, J. S. 1974. The distribution of salt pans on tidal salt marshes. Journal of Biogeography: 57–62.

Pollock, K. H, S. R Winterstein, C. M Bunck, and P. D Curtis. 1989. Survival Analysis in Telemetry Studies - the Staggered Entry Design. Journal of Wildlife Management 53: 7–15.

Pontius, Robert G., Emily Shusas, and Menzie McEachern. 2004. Detecting important categorical land changes while accounting for persistence. Agriculture, Ecosystems & Environment 101: 251–268.

Redfield, Alfred C. 1972. Development of a New England salt marsh. Ecological monographs 42: 201–237.

Rice, William R. 1989. Analyzing tables of statistical tests. Evolution 43: 223–225.

Robinson, Jeremiah. 2018. Effects of Natural and Anthropogenic Forcing on Marsh Channel Evolution.

Sallenger, Asbury H., Kara S. Doran, and Peter A. Howd. 2012. Hotspot of accelerated sea-level rise on the Atlantic coast of North America. Nature Climate Change 2. Nature Publishing Group: 884–888.

Scheiner, Sam. 1998. Design and analysis of ecological experiments. CRC Press.

Schepers, Lennert, Matthew Kirwan, Glenn Guntenspergen, and Stijn Temmerman. 2017. Spatio-temporal development of vegetation die-off in a submerging coastal marsh. Limnology and Oceanography 62. Wiley Online Library: 137–150.

Silliman, Brian R., Edwin Grosholz, and Mark D. Bertness. 2009. Human impacts on salt marshes: a global perspective. Univ of California Press.

Smith, John Bernhard. 1907. The New Jersey Salt Marsh and Its Improvement. New Jersey Agricultural Experiment Stations.

Smith, Joseph A. M., Steven F. Hafner, and Lawrence J. Niles. 2017. The impact of past management practices on tidal marsh resilience to sea level rise in the Delaware Estuary. Ocean & Coastal Management 149: 33–41.

Smith, Joseph A.M. 2013. The role of Phragmites australis in mediating inland salt marsh migration in a Mid-Atlantic estuary. PloS one 8: e65091.

Smith, Joseph A.M., and LJ Niles. 2016. Are Salt Marsh Pools Suitable Sites for Restoration? Wetland Science and Practice 33: 101–109.

Taylor, Lotem, David Curson, Gregory M. Verutes, and Chad Wilsey. 2020. Mapping sea level rise impacts to identify climate change adaptation opportunities in the Chesapeake and Delaware Bays, USA. Wetlands Ecology and Management. https://doi.org/10.1007/s11273-020-09729-w.

Theobald, David M., Don L. Stevens, Denis White, N. Scott Urquhart, Anthony R. Olsen, and John B. Norman. 2007. Using GIS to generate spatially balanced random survey designs for natural resource applications. Environmental Management 40: 134–146.

Watson, Elizabeth Burke, Cathleen Wigand, Earl W. Davey, Holly M. Andrews, Joseph Bishop, and Kenneth B. Raposa. 2016. Wetland Loss Patterns and Inundation-Productivity Relationships Prognosticate Widespread Salt Marsh Loss for Southern New England. Estuaries and Coasts: 1–20. https://doi.org/10.1007/s12237-016-0069-1.

White, G. C, and K. P Burnham. 1999. Program MARK: Survival estimation from populations of marked animals. Bird Study 46 Supplement: 120–138.

Wigand, Cathleen, Thomas Ardito, Caitlin Chaffee, Wenley Ferguson, Suzanne Paton, Kenneth Raposa, Charles Vandemoer, and Elizabeth Watson. 2017. A climate change adaptation strategy for management of coastal marsh systems. Estuaries and Coasts 40. Springer: 682–693.

Wilen, Bill O., and M. K. Bates. 1995. The US Fish and Wildlife Service’s national wetlands inventory project. Vegetation 118: 153–169.

Wilson, Carol A., Zoe J. Hughes, Duncan M. FitzGerald, Charles S. Hopkinson, Vinton Valentine, and Alexander S. Kolker. 2014. Saltmarsh pool and tidal creek morphodynamics: Dynamic equilibrium of northern latitude saltmarshes? Geomorphology 213: 99–115.

Wilson, Kristin, Joseph T. Kelley, Arie Croitoru, Michele Dionne, Daniel F. Belknap, and Robert Steneck. 2009. Stratigraphic and ecophysical characterizations of salt pools: dynamic landforms of the Webhannet salt marsh, Wells, ME, USA. Estuaries and Coasts 32: 855–870.

Wilson, Kristin, Joseph T. Kelley, Benjamin R. Tanner, and Daniel F. Belknap. 2010. Probing the Origins and Stratigraphic Signature of Salt Pools from North-Temperate Marshes in Maine, U.S.A. Journal of Coastal Research: 1007–1026. https://doi.org/10.2112/JCOASTRES-D-10-00007.1.

Yang, Zizang, Edward P. Myers, Adeline M. Wong, and Stephen Alston White. 2008. VDatum for Chesapeake Bay, Delaware Bay, and adjacent coastal water areas tidal datums, and sea surface topography.

Zeff, Marjorie L. 1988. Sedimentation in a salt marsh-tidal channel system, southern New Jersey. Marine Geology 82. Elsevier: 33–48.

Zeff, Marjorie L. 1999. Salt Marsh Tidal Channel Morphometry: Applications for Wetland Creation and Restoration. Restoration Ecology 7: 205–211. https://doi.org/10.1046/j.1526-100X.1999.72013.x.

